# TRAIL-induces Src mediated MEK/ERK, SMAD3 and β-catenin signalling in apoptosis resistant NSCLC cells

**DOI:** 10.1101/2022.08.10.503463

**Authors:** Margot de Looff, Win Sen Heng, Steven de Jong, Frank A.E. Kruyt

## Abstract

Tumour-necrosis factor related apoptosis-inducing ligand (TRAIL) receptors (TRAIL-R1 and -R2) are appealing therapeutic targets to eradicate tumours specifically via caspase-dependent apoptosis. However, resistance is often observed and TRAIL-R activation can even activate pro-tumorigenic non-canonical signalling pathways. Previously, we found that TRAIL-induced RIPK1-Src-STAT3 signalling was mediating cell migration and invasion in resistant non-small cell lung cancer (NSCLC). Here, the contribution of Src in TRAIL signalling in NSCLC cell lines was further examined. TRAIL sensitive H460 and resistant A549 NSCLC cells showed distinct time-dependent rhTRAIL-induced Src phosphorylation patterns with early activation in A549 cells. Pharmacological Src inhibition as well as shRNA knockdown or CRISPR/CAS9-dependent knockout of Src expression did not alter sensitivity to rhTRAIL-induced apoptosis in both cell lines. Silencing of secondary complex proteins showed that TRADD, but not TRAF2, FADD nor caspase-8, was required for Src activation in A549 cells. Possible mediators of Src-dependent rhTRAIL signalling were identified by Src co-IP-LC-mass spectrometric analyses. In A549 cells the number of Src-interacting proteins increased after rhTRAIL treatment, whereas protein numbers decreased in H460 cells. In rhTRAIL treated A549 cells, Src biding proteins included components of the RAF-MEK1/2-ERK, Wnt and SMAD3 signalling pathways. Functional analyses showed that Src mediated phosphorylation of MEK1/2 and ERK, prevented phosphorylation of SMAD3 and was required for nuclear translocation of ERK and β-catenin in A549 cells. Clonogenic growth of both Src proficient and deficient A549 cells was not affected by rhTRAIL exposure, although Src depletion and MEK1/2 inhibition reduced colony size and numbers significantly. In conclusion, rhTRAIL-induced and Src dependent MEK/ERK, SMAD3 and β-catenin signalling may contribute to the known pro-tumorigenic effects of rhTRAIL in resistant NSCLC cells. However, this needs to be further examined, as well as the potential therapeutic implications of targeting these pathways when combined with TRAIL receptor agonists.

## Introduction

Non-small cell lung cancer (NSCLC) is the most prevalent lung cancer type, accounting for approximately 85% of all lung cancers (1). The average 5-year survival rate is 18-20% (2). Surgery combined with (neo)adjuvant chemotherapy or radiotherapy is currently the main treatment for early stage lung cancer. Tyrosine kinase inhibitors and, more recently, immune checkpoint inhibitors have been successfully used in subgroups of patients with advanced disease, however, novel treatment strategies are still needed to improve overall prognosis of NSCLC (3–5).

Tumour necrosis factor (TNF)-related apoptosis-inducing ligand (TRAIL) receptors have been identified as promising therapeutic targets based on their tumour-selective apoptosis-inducing activity (6–8). TRAIL receptor 1 (-R1) and TRAIL-R2 induce apoptosis via their intracellular death effector domains. These domains subsequently recruit the Fas-Associated protein with Death Domain (FADD) and pro-caspase 8 to establish the death-inducing signalling complex (DISC) where caspase 8 is cleaved and activated. Subsequent effector caspase activation results in irreversible apoptosis. This pathway, also known as the death receptor/extrinsic apoptotic pathway, often involves simultaneous caspase 8-dependent cleavage of Bid, leading to cytochrome C release from mitochondria, apoptosome assembly and activation of caspase 9 and further downstream caspase activation, also known as the intrinsic/mitochondrial apoptotic pathway (9).

Various TRAIL receptor agonists have been developed and assessed. However, intrinsic and acquired resistance have been observed frequently in both *in vitro* and *in vivo* preclinical models (10,11). Therapeutic targeting of TRAIL receptors has thus far shown limited efficacy in clinical studies, although currently new studies with novel and perhaps better TRAIL formulations are under evaluation (11–14). Besides resistance towards apoptosis, TRAIL receptor activation can induce unwanted pro-tumorigenic and even metastasis-promoting effects by activation of non-apoptotic signalling pathways (14,15). These non-canonical pathways involve the formation of a secondary signalling complex consisting of the receptor-interacting serine/threonine protein kinase 1 (RIPK1), TNF receptor associated factor 2 (TRAF2), the TNF receptor type 1 associated death domain (TRADD), FADD and caspase 8 (15,16). Previously, we reported TRAIL-dependent activation of the RIPK1-Src-STAT3 pathway in resistant NSCLC cells that contributed to tumour cell migration and invasion (17).

Src is often overexpressed or hyper-activated in cancer and known to be involved in oncogenic processes like cell proliferation, survival and metastatic spread (18–20). Src comprises different functional domains, including two Src homology (SH) domains and a catalytic domain. Its activity is mainly regulated by two phosphorylation sites, the positive regulatory auto-phosphorylation at Tyr418 in the catalytic domain and the negative regulatory Tyr530 phosphorylation at the C-terminal part (21,22). Elevated Src expression and activation has been reported in the majority of lung cancers, especially in NSCLC (20,23). Activated Src can phosphorylate and thereby inactivate caspase 8 resulting in TRAIL resistance (24,25). Furthermore, Src activation has been implicated in Akt activation after TRAIL treatment in breast cancer cells and inhibition of Src sensitized hepatocellular carcinoma cells for TRAIL-induced apoptosis (26,27). In the current study, we examined the possible role of Src and underlying mechanisms in modulating apoptotic and pro-tumorigenic TRAIL signalling in NSCLC cells.

## Material and methods

### Cell lines

A549 and H460 cells were obtained from the ATCC and cultured in RPMI 1640 (Gibco, Waltham, USA) with 10% FBS (Bodinco, Alkmaar, The Netherlands). Cells were maintained in a humidified 5% CO2 atmosphere at 37°C. The cell lines were tested annually for authenticity by short tandem repeat profiling DNA fingerprinting (Baseclear, Leiden, The Netherlands) and for mycoplasm by PCR.

### Reagents

Human rhTRAIL was obtained from Peprotech Inc. (London, UK). Pharmacological inhibiters used were Dasatinib (Axon Medchem, Groningen, The Netherlands), Selumetinib (AZD6244, Axon Medchem) and SIS3 (Cayman chemical, Ellsworth, USA).

### Modulation of Src expression

Short interfering (si)RNAs against TRADD (SR305738 Origene, Rockville, USA), FADD (SR305777 Origene), TRAF2 (SR304927 Origene) and caspase 8 (sc-29930 Santa Cruz, Dallas, USA) were transfected in cells with Oligofectamine reagent (ThermoFisher, Waltham, USA) according manufactures protocol. In short, 3.5*10^5^ cells were plated in a 6 wells plate and incubated overnight. Subsequently, cells were washed with PBS and 800 µl optimum medium (Gibco) was added, prior to adding short interfering (si)RNA transfection mix. Transfection mix was prepared: 3 µl Oligofectamine was dissolved in 12 µl Opti-MEM per well and incubated for 10 min at RT. Next, 185 µl with 1 µM siRNA was added and the transfection mix was incubated for 20 min at RT. In each well 200 µl transfection mix was added dropwise to the cells and incubated for 4 hrs at 37°C followed by addition of 500 µl medium with 30% FCS. Short hairpin RNA silenced Src in A549 cells were described previously (17). Src gene knockout (A549-Src KO) and empty vector control (A549-Src ctrl) cells were generated by CRISPR-Cas9 technology. crRNAs were designed using https://benchling.com. DNA oligonucleotides for Src exon 4: GTCCTTCAAGAAAGGCGAG (guide 1) and exon 5: AGCCCAAGGATGCCAGCCAG (guide 2) were ordered from IDT (Leuven, Belgium) and cloned into pSpCas9(BB)-2A-GFP(PX458) (Addgene Teddington, UK), according to the protocol of Ann Ran *et al*. (28). After transformation in bacteria (One Shot™ TOP10 Chemically Competent E. coli; Thermo Fisher Scientific, Bleiswijk, Netherlands), successful cloning was validated by sequencing. The Src KO and empty vector (Src ctrl) constructs were transfected in A549 cells with a FuGENE® HD reagent-DNA ratio of 3:1 according to manufacturer’s protocol. Briefly, 2,5*10^5^ A549 cells per well (6-well plate) were incubated overnight for transfection. After 48 hrs of transfection, GFP positive cells were single cell sorted with the MoFlo cell sorter (Beckman Coulter, Brea, USA). Clonal cultures were evaluated for A549-Src KO by western blot and a representative clone was selected.

### MTT cell viability assay

1*10^4^ cells in 100 µl medium per well were plated in a 96-wells plate (Greiner Bio-One, Alphen aan den Rijn, The Netherlands) and incubated overnight. 100 µl medium with or without rhTRAIL was added for 24 or 48 hrs yielding a total volume of 200 µl per well. Next, 20 µl of 5 mg/ml Thiazolyl Blue Tetrazolium Bromide (MTT; Sigma-Aldrich, Saint Louis, USA) solution in PBS was added per well and incubated at 37°C for 3 hrs. The plates were centrifuged at 900 rpm for 15 min without brake. The formazan crystals were dissolved using 200 µl dimethyl sulfoxide (DMSO; Merck, Burlington, USA) and absorbance was measured at 520 nm (Biorad, Hercules, USA).

### Western blot analysis

Cells, treated as indicated, were washed twice with ice-cold PBS and lysed with M-Per (ThermoFisher) including 100x Halt phosphatase and protease inhibitors (ThermoScientific, Waltham, USA) for 1.15 hrs and centrifuged at 14.000 rpm at 4°C for 10 min. Protein concentrations were determined by Bradford protein assay (29). 20 μg of protein per sample was loaded and separated on 8-12% SDS–PAGE gels and electro blotted onto polyvinylidenedifluoride (PVDF) membranes (Immobilon-P PVDF membrane 0.45μm; Merck-Millipore, Burlington, USA). Subsequently, membranes were blocked for 1 hr at RT in 5% albumine bovine fraction V (Thermo Scientific, pH 7.0) washed in TBS + 0.05% Tween with pH 8.0 (TBS-T) and incubated overnight at 4°C with the primary antibodies diluted in TBS-T. The membranes were washed thrice for 5 min in TBS-T and incubated with Horseradish peroxidase secondary antibodies (DAKO, Santa Clara, USA) for 1 hr at RT. After incubation, the membrane was washed twice for 5 min in TBS-T, followed by 5 min washing in TBS. Subsequently the blots were incubated with the chemiluminescent Lumi-Light (Roche, Basel, Switzerland) and bands visualized with the Chemidoc imaging system (BioRad). The following primary antibodies were used. From Cell Signaling Technologies (Danvers, USA): Src (#2109), pSrc tyr416 (#2101), pSrc tyr527 (#2105), Caspase 8 (#9746), TRADD (#3684), TRAF2 (#4724), FADD (#2782), Erk1/2 (#9102), pErk thr202/tyr204 (#9106), MEK1 (#2352), MEK2 (#9147), pMEK1/2 Ser217/221 (#9154), SMAD3 (#9523) and Lamin A/C (#4777). Other antibodies used: β-actin (MP biomedicals), GAPDH (Ab128915; Abcam, Cambridge, UK), pSMAD3 (Ab52903, Abcam) and β-catenin (BD biosciences, Franklin Lakes, USA). All antibodies used were dissolved in 5% albumine bovine fraction V (Thermo Scientific, pH 7.0) in TBS-T.

### Immunoprecipitation

The cells were grown in a T165 flask until 60-70% confluency followed by treatment as indicated. The cells were washed twice with ice cold PBS and scraped in cold NP-40 buffer (50 mM Tris pH 7.7, 150 mM NaCl and 0.5% v/v Igepal) containing 100x Halt phosphatase and protease inhibitors (ThermoFisher). Bradford assay was performed to determine protein concentration and samples were stored on ice, or at −20°C until use. A beads-antibody complex was prepared prior the actual immunoprecipitation. For each condition: 1,5 mg of superparamagnetic Dynabeads protein G (ThermoFisher) were isolated and 10 µg Src antibody (clone GD11; Sigma Aldrich) diluted in 200 µl PBS + 0.1% Tween-20 (PBS-T) (#P1379; Sigma-Aldrich) was added. The beads-antibody solution was rotated for 10 min at RT, and washed once in 200 µl PBS-T. Subsequently, the complex was washed twice in 200 µl conjugation Buffer (20 mM Sodium Phosphate, 0.15M NaCl pH 7-9), resuspended in 250 µl 5 mM BS^3^ (Life technologies, Carlsbad, USA) and incubated for 60 min at RT with rotation. The cross-linking reaction was quenched by adding 12.5 µl quenching buffer (1M Tris HCl pH 7.5) for 15 min with rotation at RT. The Src antibody-beads complexes were washed thrice with 200 µl PBS-T and 1.5 mg antibody-beads were mixed with 500 µg protein per sample. Binding of the beads-antibody-protein complex was allowed for 1 hr under rotation at 4°C. The beads-antibody-protein complexes were washed thrice with 200 µl washing buffer (citrate-phosphate buffer, pH 5.0) and the beads-antibody complexes were mixed with 20µl elution buffer (0.1 M citrate; pH 2) including 6x SDS loading buffer. The mixture was heated for 10 min at 70°C and proteins were separated by 8-12% SDS page gel for western blot analyses.

### Mass spectrometry

Src was co-immunoprecipitated as described above in control and 1 hr rhTRAIL exposed A549 and H460 cells. A549-Src-KO cells served as a non-specific binding control The protein mixtures were loaded on a RunBlue 1 mm*10 well 8%-Bis-Tris – gel (Expedeon, Cambridge, UK) after heating the mixture at 70°C for 10 min. Whole gel processing procedure was performed as described previously (30). LC-MS/MS was performed by the Ultimate 3000 HPLC system (Thermo Scientific) coupled online to a Q-Exactive-Plus mass spectrometer with a NanoFlex source (Thermo Scientific).

### Data processing

The mass spectrometry data was processed as described previously (31). In short, the PEAKS 8.0 (Bioinformatics Solutions Inc., Waterloo, Ontario, Canada) software was applied to the spectra generated by the Q-exactive plus mass spectrometer to search against a Human Protein database (SwissProt containing 20197 entries) using fixed modification carbamidomethylation of cysteine and the variable post translational modifications oxidation of methionine with a maximum of 5 posttranslational modifications per peptide at a parent mass error tolerance of 10 ppm and a fragment mass tolerance of 0.03 Da. False discovery rate was set at 0.1% and at least 2 unique peptides per protein should be present. Proteins were corrected for background/contamination by eliminating proteins present in IgG1, IgG2b and A549-Src KO-Src IP samples and the fold change (FC) and Log2(FC) of the remaining proteins were calculated. To identify proteins of interest, a STRING network analysis was performed as well as a KEGG pathways analysis.

### Immunofluorescence microscopy

In 96 wells plates 5000 cells per well were seeded and incubated overnight. The cells were treated as indicated and fixed with 100 µl 2% paraformaldehyde for 15 min. Subsequently, cells were washed twice with PBS, permeabilized with PBS + 0.5% Tween for 10 min, washed twice with 100 µl PBS + 0.5% Tween, blocked for 1 hr in 2% BSA + 0.1% Tween20 + 1:50 normal goat serum (DAKO) in PBS and washed twice with PBS + 0.5% Tween. The primary antibodies diluted in blocking buffer were added and incubated for 1 hr. Primary antibodies used were: Src (#2109; Cell Signaling Technologies), β-catenin (BD biosciences), Mouse IgG (#610153; BD biosciences) and Rabbit IgG (Cell Signaling Technologies). Secondary antibodies: Alexa488 IgG (Invitrogen), Alexa 568 (Invitrogen). After washing twice with 100 µl PBS + 0.5% Tween the secondary antibody diluted 1:200 in blocking buffer was added and incubated for 1 hr. Subsequently, cells were washed twice with PBS + 0.5% Tween followed by incubation with 2 mg/ml DAPI (Sigma) for 10 min, washed twice with PBS and stored at 4°C until analysis. Pictures were taken with the EVOS digital colour fluorescence microscope (Invitrogen, Carlsbad, USA) and analysed.

### Clonogenic assay

For colony formation, 200 cells per well were seeded in a 6 wells plate and rhTRAIL (50 ng/ml), selumetinib (Selu; 0.1 µM) or SIS3 (3 µM) were added after overnight incubation. After 10 days cells were fixed with methanol for 15 min, stained with 0.1% crystal violet solution and washed thoroughly with water and plates were dried. Colony numbers and size were analysed using the Vspot spectrum (AID, Strasbourg, Germany) and the colony counter plugin from ImageJ with a minimum-maximum size (Pixel^2^) 50-100000, circularity 0.8-1 and a minimum area of 20.

### Statistics

All experiments were performed at least 3 times independently, unless otherwise indicated.

## Results

### Src does not modulate rhTRAIL induced apoptotic signalling in A549 and H460 cells

TRAIL resistant A549 and sensitive H460 cells were examined for Src activation after rhTRAIL (50 ng/ml) exposure for various time periods. Different phosphorylation patterns of Src-Y418p (activated Src) were observed in A549 and H460 cells. Src-Y418p was detected at early timepoints in A549 but not in H460 cells (Fig. 1A). In H460 cells very low basal levels of Src-Y418p were detected with no increases within 60 min of rhTRAIL treatment, whereas high levels of Src-Y418p were detected at later time points that were not seen in A549 cells (Fig. 1A). To investigate whether Src regulates rhTRAIL sensitivity, Src activity was inhibited by Dasatinib or its expression downregulated by either shRNA-dependent silencing (knockdown, KD) or genetic ablation (knockout, KO) using CRISPR/Cas9 gene editing. Dasatinib effectively prevented phosphorylation of Src-Y418p in A549 and H460 cells and Src expression was effectively silenced in A549-Src KD cells and depleted in A549-Src KO cells compared to A549-Src ctrl (Fig. 1B-C). Src inhibition did not significantly affect rhTRAIL sensitivity in rhTRAIL resistant or sensitive cells (Fig. 1D). Also A549 Src-KD and -KO cells showed no altered rhTRAIL sensitivity as compared to parental A549 and A549-Src ctrl cells (Fig. 1E). Overall, rhTRAIL treatment resulted in differential Src activation in sensitive and resistant NSCLC cells. However, Src was not instrumental for rhTRAIL-mediated apoptosis.

**Figure 1.**
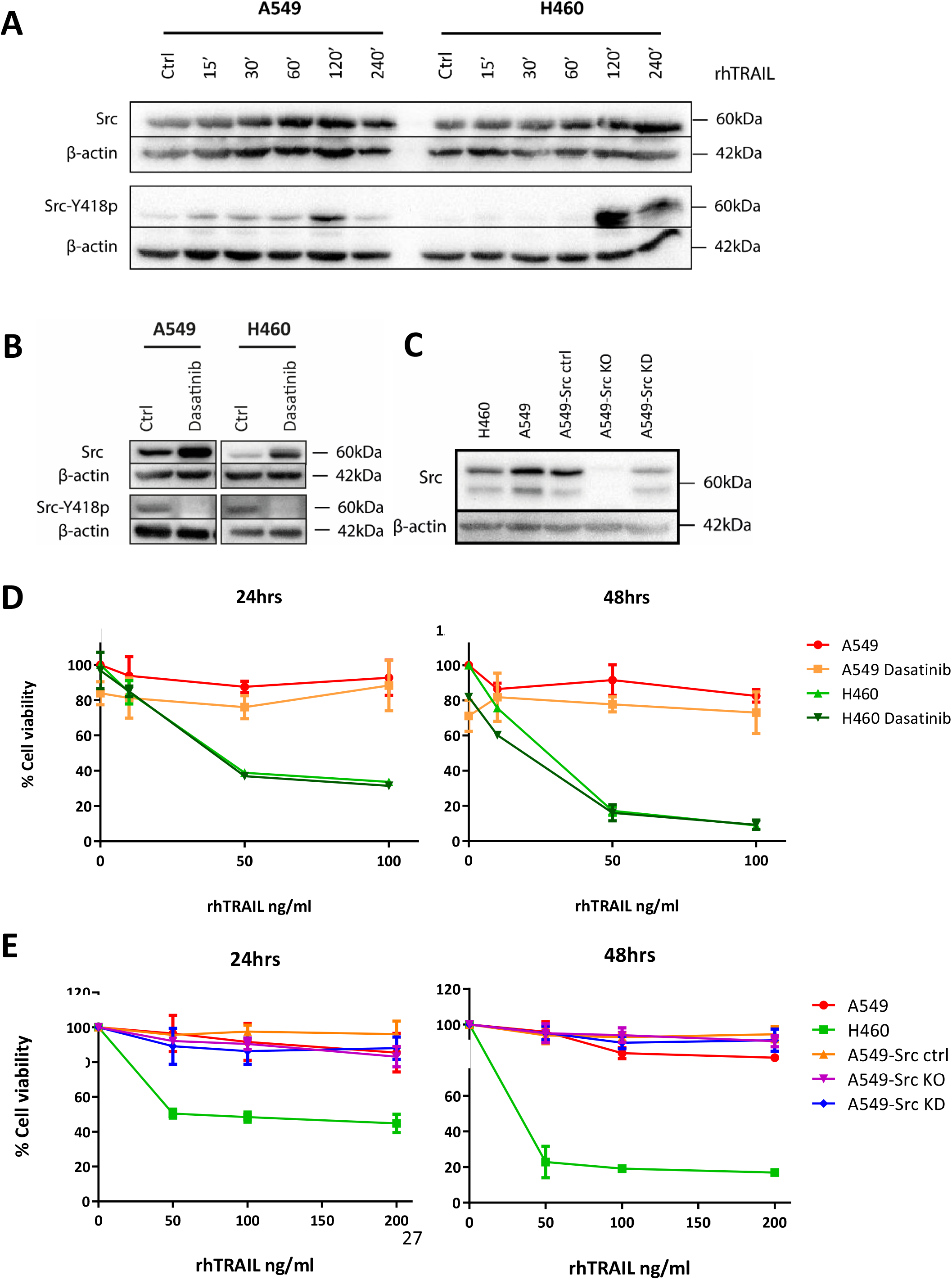
Src is differentially activated in A549 and H460 cells and does not regulate TRAIL induced apoptosis. **(A)** Western blot analysis of total and phosphorylated Src in A549 and H460 cells after treatment with 50 ng/ml rhTRAIL for the indicated time periods. In resistant A549 cells Src phosphorylation was detected at early time points. In sensitive H460 cells, Src phosphorylation was detected only at later time points. **(B)** Western blot analysis of total Src and Src-Y418p showing potent Src inhibition after treatment with 1 µM Dasatinib for 24 hrs. **(C)** Src expression levels in H460, A549, A549-Src ctrl, CRISPR/CAS9 knockout (A549-Src KO) and short hairpin knockdown (A549-Src KD). **(D)** MTT assays measuring cell viability of A549 and H460 cells treated with different concentrations rhTRAIL and in absence or presence of Dasatinib for 24 or 48 hrs. **(E)** MTT assays of H460, A549, A549-Src ctrl, A549-Src KD and A549-Src KO cells treated with indicated concentrations rhTRAIL for 24 or 48 hrs. Neither inhibition nor KD or KO of Src affected sensitivity towards rhTRAIL induced apoptosis. MTT results are expressed as mean ± SD of three independent assays.

### TRADD is required for rhTRAIL mediated Src activation

Previously we demonstrated that TRAIL-induced Src activation is mediated by RIPK1 (32). To further examine the involvement of other DISC and secondary complex components, caspase 8, FADD, TRADD and TRAF2 were silenced with specific siRNAs in A549 cells. Caspase 8, FADD, and TRAF2 silencing did not alter Src activation conclusively (data not shown). However, TRADD silencing delayed Src activation when compared to control cells and resulted in increased caspase 8 cleavage (Fig. 2). Thus, TRADD mediated rhTRAIL-induced Src activation and suppressed caspase 8 cleavage.

**Figure 2.**
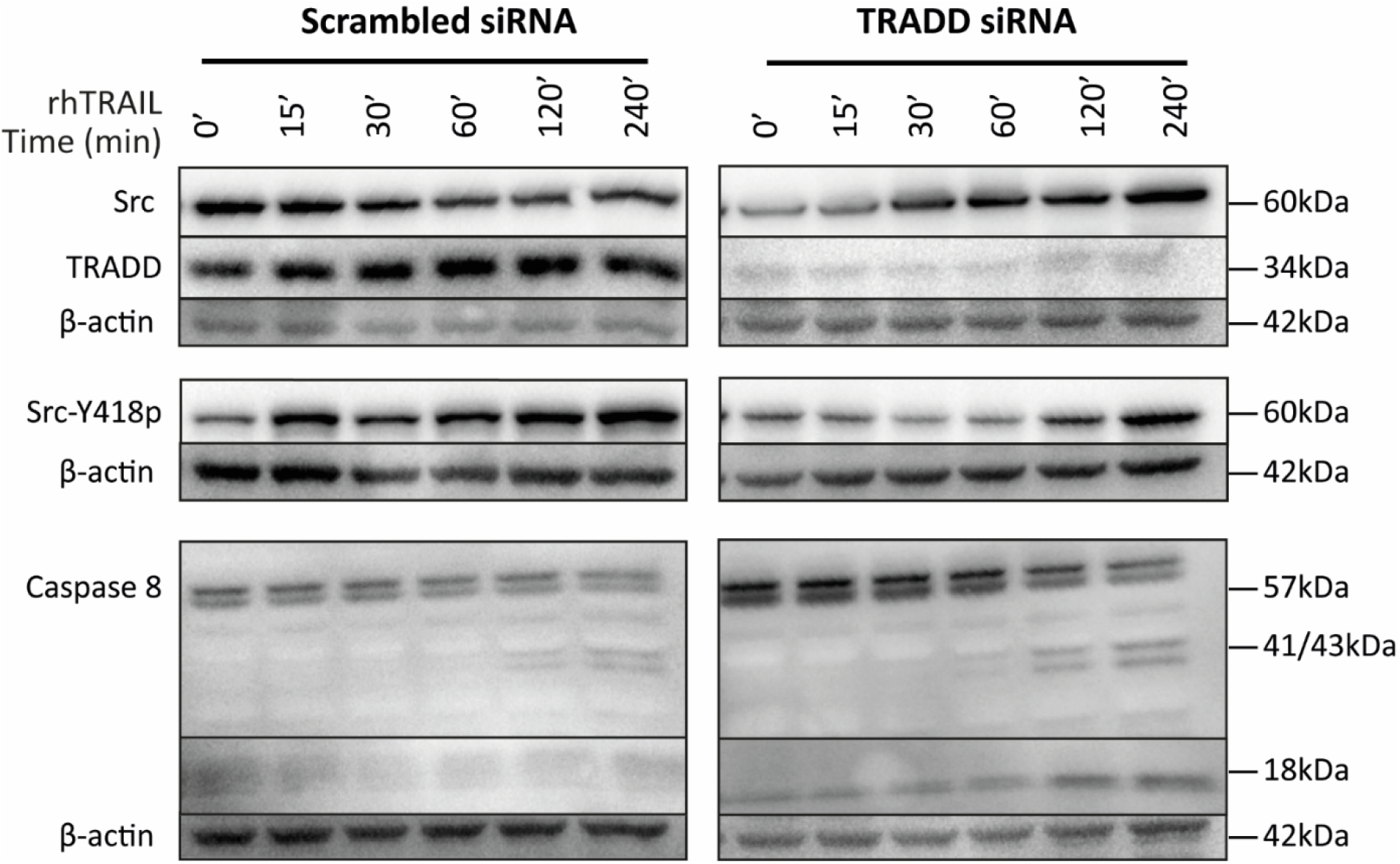
TRADD mediates rhTRAIL-induced Src activation in A549 cells. Protein levels of Src, TRADD, Src-Y418p and caspase 8 in rhTRAIL treated A549 cells transfected with control scrambled siRNA or TRADD siRNA were examined by western blotting, indicating that TRADD is involved in Src activation at early time points.

### Analysis of the Src protein interactome

To further elucidate the underlying molecular mechanisms of Src-mediated pro-tumorigenic rhTRAIL signalling, we set out to study the interactome of Src in A549 and H460 cells. Cells were treated for 1 hr with 50 ng/ml rhTRAIL, followed by Src immunoprecipitation and tryptic peptide-based mass-spectrometry to identify possible differential interacting proteins. In untreated A549 cells 314 proteins were found to interact with Src, increasing to a total of 435 proteins after rhTRAIL treatment, of which 181 were newly interacting proteins (Fig. 3A, protein lists in Supplementary data 1). On the contrary, in H460 cells rhTRAIL treatment resulted in a reduction of Src binding proteins, from 560 proteins to 180 proteins, respectively, with only 21 newly bound proteins (Fig. 3A, protein lists in Supplementary data 1). In both untreated cell lines, 220 overlapping Src-interacting proteins were found, and 125 overlapping proteins in rhTRAIL treated cells. In rhTRAIL exposed A549 cells 87 unique proteins interacted with Src, compared to only 11 unique proteins in rhTRAIL treated H460 cells. Taken together, substantially differences in the number of Src binding proteins and interactome composition between H460 and A549 cells were found in both untreated and rhTRAIL-treated conditions. Notably, an overall increase in the number of interacting proteins and detection of unique interactors was seen in rhTRAIL-treated A549 cells, whereas in H460 cells protein numbers decreased and only a small number of unique interactors were detected.

**Figure 3.**
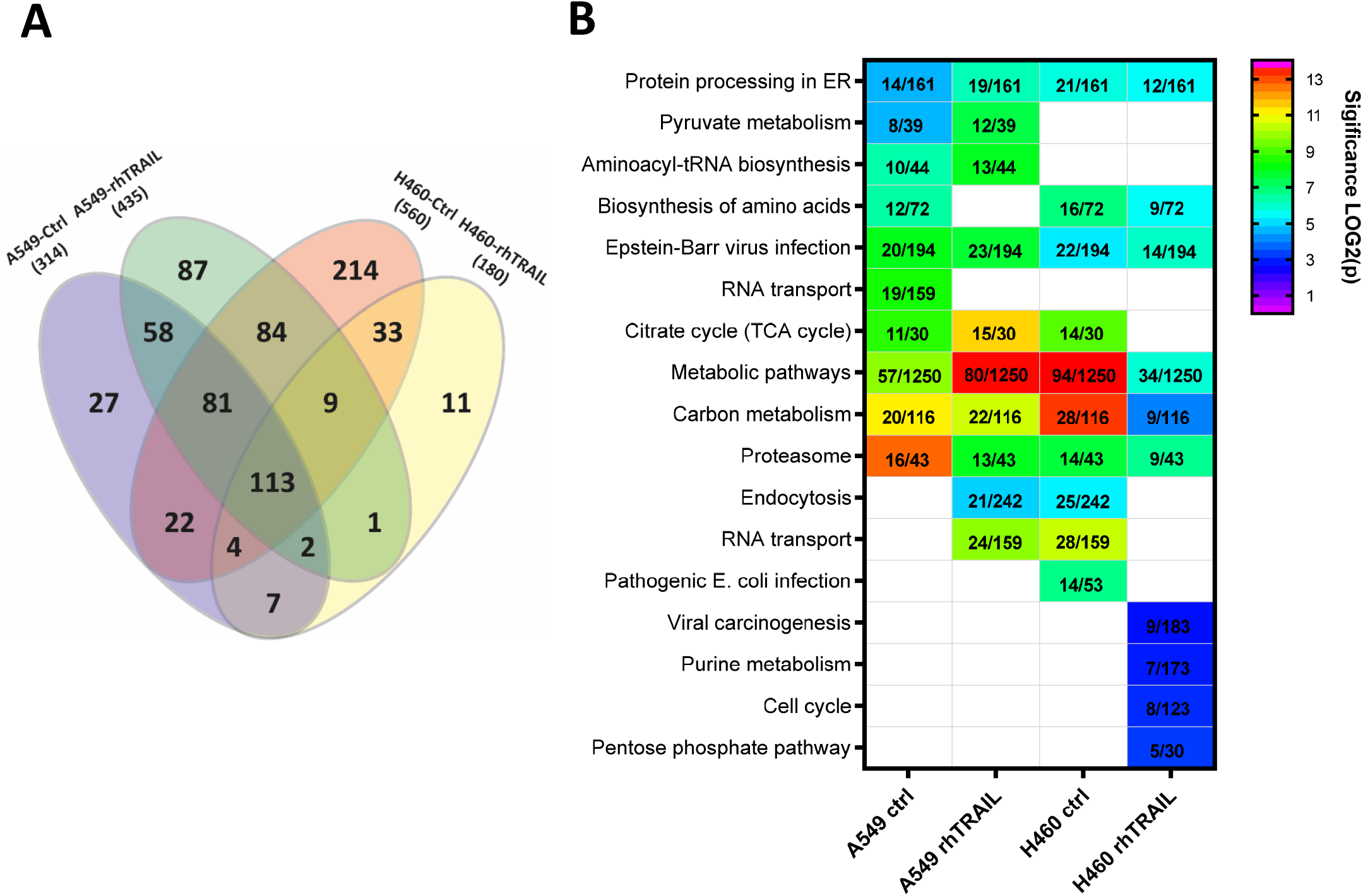
Analyses of Src interactomes in H460 and A549 cells with(out) rhTRAIL treatment. Src co-IP was performed in H460 and A549 cells −/+ rhTRAIL for 1 hr followed by mass-spectrometry (MS) analyses to examine the Src interactomes (n=1). **(A)** Venn diagrams representing proteins found in the Src interactomes of A549-ctrl, A549-rhTRAIL, H460-ctrl and H460-rhTRAIL cells. After rhTRAIL treatment, the number of proteins in the Src interactome of A549 increased and of H460 decreased. **(B)** Heatmap showing the top 10 most significantly represented KEGG pathways based on analyses of the Src interactomes of A549-ctrl, A549-rhTRAIL, H460-ctrl and H460-rhTRAIL cells. Significant p values from the false discovery rate are depicted with their -log10(p) values by a colour range. The number of proteins present in the interactomes per KEGG pathway is shown.

KEGG pathway analyses were performed subsequently with all Src interacting proteins to obtain insight in associated biological processes (Fig. 3B). Top 10 significant annotations were highly similar between the A549 and A549-rhTRAIL Src interactomes, representing predominantly metabolic and biosynthesis pathways, with endocytotic pathways being unique in rhTRAIL treated cells (Fig. 3B). In the H460 and H460-rhTRAIL Src interactomes 4 out of 10 pathways overlapped and most processes were linked with nucleotide and metabolic pathways (Fig. 3B). Opposed to the A549 interactomes, proteins involved in endocytosis were found in complex with Src in untreated H460 cells and that were not seen in H460-rhTRAIL interactomes. Next, proteins showing at least a 2-fold increased or decreased binding to Src after rhTRAIL exposure were selected for further evaluation. In A549 cells interactomes 182 proteins increased and 60 decreased, whereas in H460 cells 21 proteins increased and 60 proteins decreased (Fig. 4A; protein lists in supplementary data 1). KEGG pathway analysis revealed that proteins belonging to various pathways, including metabolism and endocytosis, were increased in A549-rhTRAIL and decreased in H460-rhTRAIL Src interactomes (Fig. 4B).

**Figure 4.**
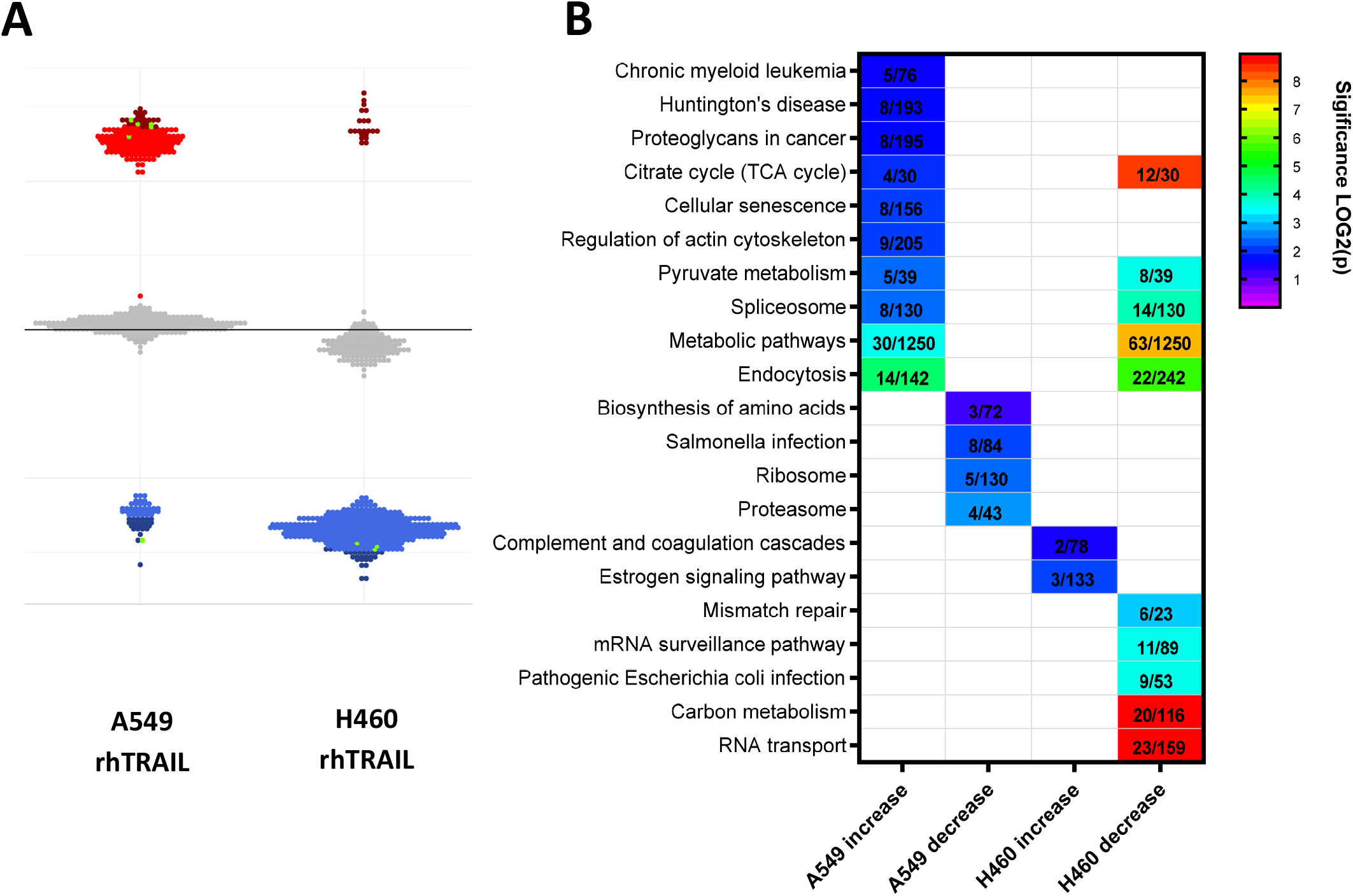
Proteins in the Src interactomes that change 2-fold or more in H460 and A549 cells with(out) rhTRAIL treatment. **(A)** Dot plots showing proteins in the Src interactomes of A549-rhTRAIL and H460-rhTRAIL. Each dot represents a protein with an at least a 2-fold change in abundancy in the corresponding interactome after rhTRAIL exposure; blue (decrease), red (increase) and grey (less than 2-fold change). The proteins indicated in dark blue or red represent the top 15% most fold-changed proteins **(B)** KEGG pathway heatmaps based on the top 15% most differential proteins detected in the Src interactomes (p=0.05) indicated in **(A)**. Significant p values from the false discovery rate are depicted with their LOG2 values by colour. The numbers of proteins representing a KEGG pathway are indicated.

To identify the potential interactors that might mediate rhTRAIL pro-tumorigenic signalling, STRING network and KEGG pathway analyses were performed on the top 15% proteins that either increased or decreased at least 2-fold in interactomes obtained after rhTRAIL treatment (4A). This selection yielded 75 upregulated proteins and 60 downregulated proteins in A549-rhTRAIL, and 21 upregulated proteins and 87 downregulated proteins in H460-rhTRAIL interactomes (protein lists supplementary data 1). Using string network analysis, in the Src interactome from rhTRAIL treated A549 cells, among others MEK1, MEK2, SMAD3 and β-catenin (Catenin-β1/CTNNB1) were upregulated (Fig. 5A; Supplementary data 1). On the other hand, MEK2, PP2A, β-catenin and Catenin-α1 were downregulated in the H460-rhTRAIL interactome (Fig. 5A; Supplementary data 1). Notably, the phosphatase PP2A was downregulated in both the A549-rhTRAIL and H460-rhTRAIL interactome, PP2A is a major Src inhibitor as well as a regulator of the Wnt signalling pathway by dephosphorylating Wnt pathway components, which can result in both suppression and stimulation of tumour growth, depending on the cellular context (Fig. 5A; Supplementary data 1) (33,34).

**Figure 5.**
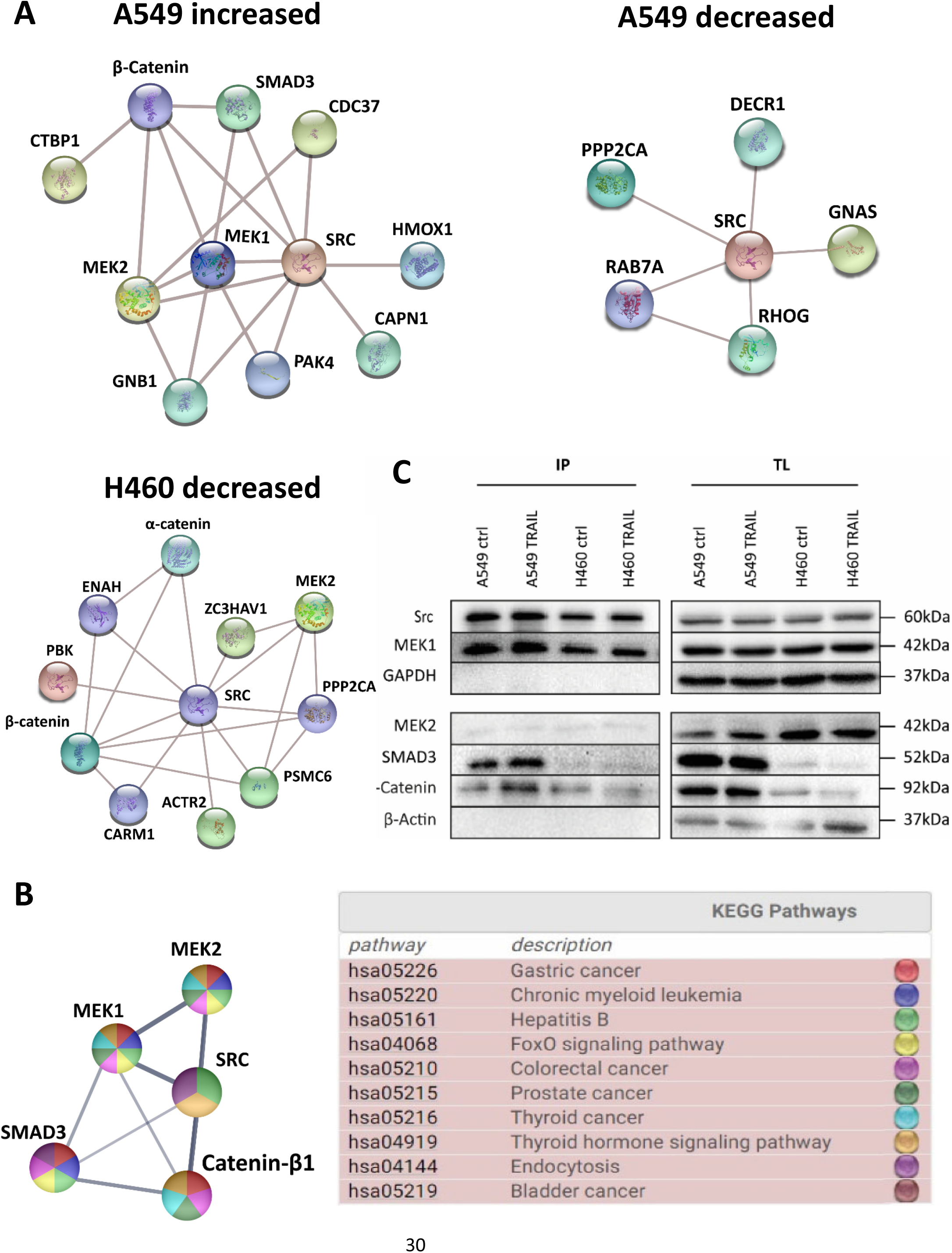
Potential TRAIL-regulated mediators of Src-dependent pro-tumorigenic signalling in A549 cells. String network analysis of the proteins identified in the Src interactomes that **(A)** increased in A549-rhTRAIL, decreased in A549-rhTRAIL, decreased in H460-rhTRAIL. **(B)** The Src-MEK1-MEK2-SMAD3-β-catenin network and KEGG pathway analyses; top 10 most significant cancer associated pathways are shown. **(C)** Src co-IP experiments to determine interactions with MEK1, MEK2, SMAD3 and β-catenin in A549 cells treatment for 1 hr with 50 ng/ml rhTRAIL. Western blots confirmed interactions between Src and MEK1, SMAD3 and β-catenin. Total lysate (TL).

KEGG pathway analysis further confirmed that Src, MEK1, MEK2, SMAD3 and β-catenin were implicated in various cancer-related pathways, being able to interact with Src as well as with each other (Fig. 5B). Such interactions might mediate Src-dependent pro-tumorigenic effects of rhTRAIL exposed A549 cells. MEK1 and MEK2 are part of the RAS mediated RAF-MEK1/2-ERK proliferation and survival pathway, which is often hyperactivated in lung cancers (35,36). SMAD3 is a mediator of TGF-β signalling and has both tumour suppressive and oncogenic functions (37). Wnt-β-catenin signalling promotes stemness, tumorigenesis and cancer cell proliferation in various cancers including NSCLC (38–40).

### MEK1/2, SMAD3 and β-catenin as possible mediators of rhTRAIL-induced Src signalling

The interactions of MEK1, MEK2, SMAD3 and β-catenin with Src were confirmed by direct co-IP/western blotting experiments (Fig. 5C). High levels of MEK1 associated with Src were detected in A549 cells independent of rhTRAIL treatment. MEK2 levels, however, were very low in both cell lines (Fig. 5C). An apparent increase in SMAD3 and β-catenin binding to Src was observed in A549-rhTRAIL cells. In contrast, β-catenin interactions decreased in rhTRAIL-treated H460 cells (Fig. 5C).

To further investigate the involvement of these proteins in Src mediated signalling, possible time dependent effects of rhTRAIL on phosphorylation of MEK1/2, ERK (that is activated by MEK1/2) and SMAD3 (Ser423/425), and the expression of β-catenin were determined in A549, A549-Src ctrl and A549-Src KO cells (Fig. 6A). Basal MEK1/2 phosphorylation levels were higher in A549 and A549-Src ctrl cells compared to Src depleted cells, whereas total MEK1 and MEK2 levels remained mostly unaltered (Fig. 6A). Phospho-MEK1/2 (pMEK1/2) levels increased after 15 min rhTRAIL, particularly in Src proficient cells. Normally, pMEK1/2 phosphorylates and activates Erk which is subsequently translocated to the nucleus. pERK is therefore a functional readout for MEK1/2 activation. After 15 mins of rhTRAIL treatment downstream ERK phosphorylation was also stronger in A549 and A549-Src ctrl cells when compared to A549-Src KO cells, while total ERK levels were unaffected (Fig. 6A). β-catenin expression slightly decreased upon rhTRAIL treatment in A549 and A549-Src ctrl cells, whereas lower basal levels were found in Src deficient cells and no clear effect of rhTRAIL on expression was seen (Fig. 6A). Levels of phospho-SMAD3 (pSMAD; Ser423/425) increased strongly upon rhTRAIL treatment in A549-Src KO cells compared to the Src proficient cells (Fig. 6A). Taken together, Src was found to be involved in rhTRAIL-induced MEK1/2 and ERK activation, the regulation of β-catenin protein levels and the suppression of SMAD3-Ser243/425 phosphorylation.

**Figure 6.**
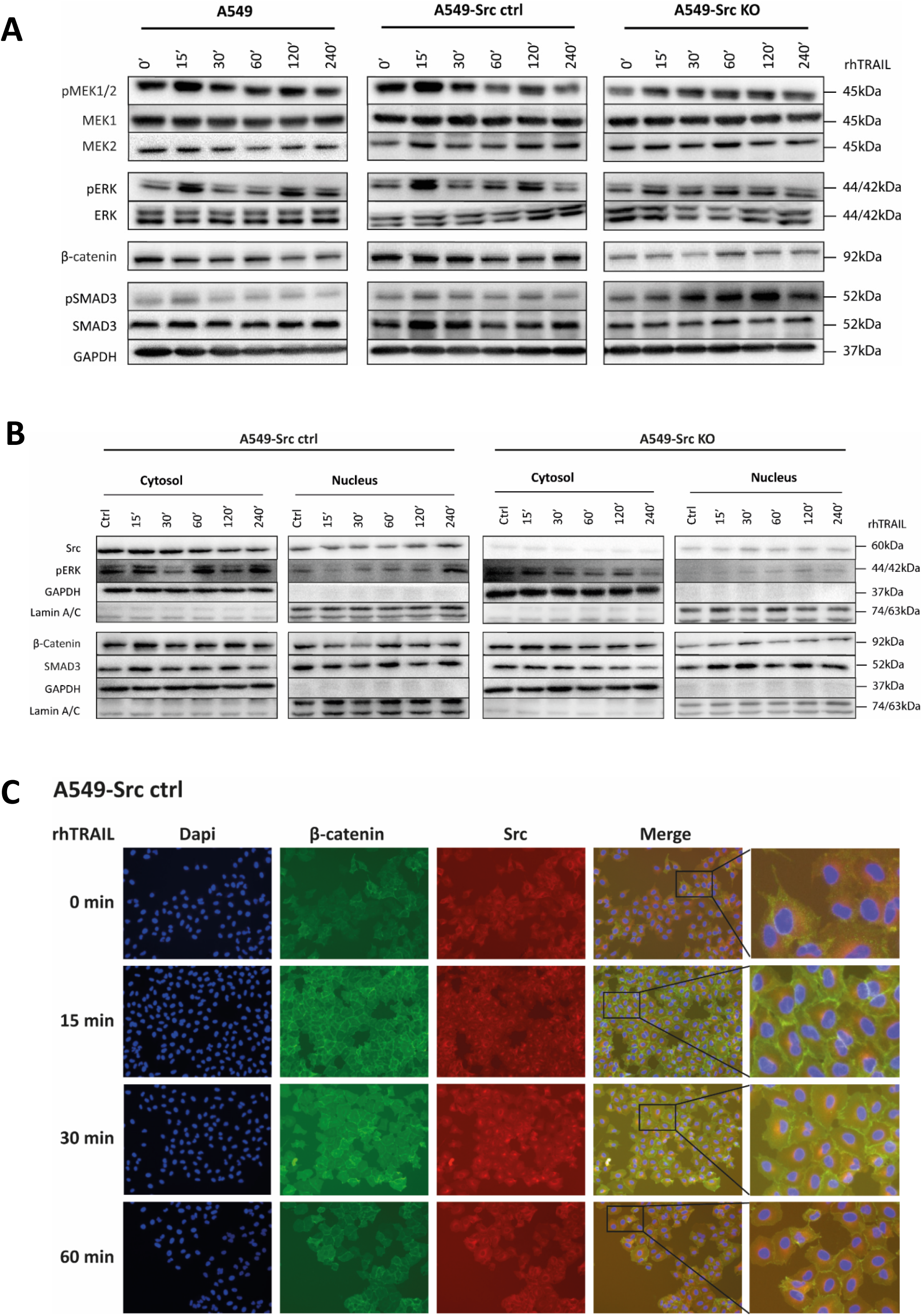
Effect of TRAIL and Src on MEK1/2, ERK, β-catenin and SMAD3 activity in A549 cells. **(A)** Western blot analysis showing expression of the indicated (phosphorylated) proteins that were identified as possible mediators of TRAIL-induced Src signalling in A549 cells. Protein expression was determined in A549, A549-Src ctrl and A549-Src KO cells treated for indicated time periods with 50 ng/ml rhTRAIL. Src appeared to be involved in the activation of MEK1/2 and ERK, regulation of β-catenin protein levels and the suppression of SMAD30-Ser243/425 phosphorylation. **(B)** Western blot analysis of proteins in cytosolic and nuclear extracts from A549-Src ctrl and A549-Src KO cells after treatment with 50 ng/ml rhTRAIL for the indicated time periods. Src appeared to be involved in TRAIL-induced pERK and β-catenin nuclear translocation. **(C)** Immunofluorescent microscopic analyses (40x) of Src and β-catenin in A549-Src ctrl cells after rhTRAIL exposure at the indicated times. DAPI (blue), β-catenin (green) and Src (red) and merged pictures and enlargements. Demonstrated here is that rhTRAIL treatment increased Src and β-catenin co-localisation in spots adjacent to the nucleus (peri-nuclear), as well as rhTRAIL dependent enhanced localisation of β-catenin to the plasma membrane.

### Effect of Src and rhTRAIL on the subcellular localization of pERK, SMAD3 and β-catenin

Next, the effects of Src on the cytoplasmic and nuclear localization of the identified downstream proteins were examined by subcellular fractionation of cell lysates and western blotting. rhTRAIL treatment in A549 cells resulted in increased Src levels in the nuclear fraction at later timepoints, concomitant with a decrease in cytoplasmic Src (Fig. 6B; Supplementary fig. 2A). pERK levels increased in the cytosol at early timepoints post rhTRAIL treatment and localized also to the nucleus after 240 min in Src proficient cells, whereas pERK levels overall decreased in Src deficient cells (Fig. 6B, supplementary fig. 2A). Total SMAD3 levels increased in the nuclear fraction of Src deficient cells after rhTRAIL treatment, whereas levels remained mostly constant in Src proficient cells (Fig. 6B; Supplementary fig. 2A). Unfortunately, we were not able to detect pSMAD3 in the nuclear fraction after subcellular fractionation. However, we did found an increase in pSMAD (Ser423/425) levels in the total lysates of Src deficient A549 cells after rhTRAIL treatment (Fig. 6A). In general, pSMAD3 is known to translocate to the nucleus, suggesting that nuclear SMAD3 represents pSMAD3 (41,42). However, this needs to be further substantiated. Nuclear β-catenin levels increased upon rhTRAIL treatment in Src proficient cells, which was less detectable in Src deficient cells (Fig. 6B; Supplementary fig. 2A).

Next, we explored Src and β-catenin localization following rhTRAIL treatment by immunofluorescent microscopy in A549 cells. We found increased co-localisation of Src and β-catenin at the periphery of the nucleus upon rhTRAIL treatment (Fig. 6C, supplementary fig. 2B). Furthermore, an increase in membrane localised β-catenin was seen upon rhTRAIL treatment, that was not observed in Src deficient cells (Fig. 6C; Supplementary fig. 2B). Together, these results showed that Src is required for rhTRAIL induced pERK nuclear translocation and a peri-nuclear localisation of β-catenin in A549 cells.

### Effects of rhTRAIL, Src, MEK1/2 and SMAD3 on clonogenic growth of A549 cells

Finally, we investigated the possible effects of Src and the identified downstream proteins MEK1/2 and SMAD3 on clonogenic growth of A549 cells (Figure 7). Src deficiency resulted in significantly reduced average colony size, but had no effect on the number of colonies (Fig. 7). Furthermore, rhTRAIL treatment did not affect colony formation or growth, neither in Src proficient nor deficient cells. SMAD3 inhibition with inhibitor of SMAD3 (SIS3) had no significant effect on colony formation and growth, whereas MEK1/2 inhibition with Selumetinib significantly reduced colony growth although independent of Src status or rhTRAIL treatment (Fig. 7B). In Src KO cells, MEK1/2 inhibition also significantly reduced the number of colonies (Fig. 7A). Thus, independent of rhTRAIL exposure, Src and MEK1/2 both stimulated colony growth in A549 cells, and MEK1/2 also enhanced colony formation in Src KO cells.

**Figure 7.**
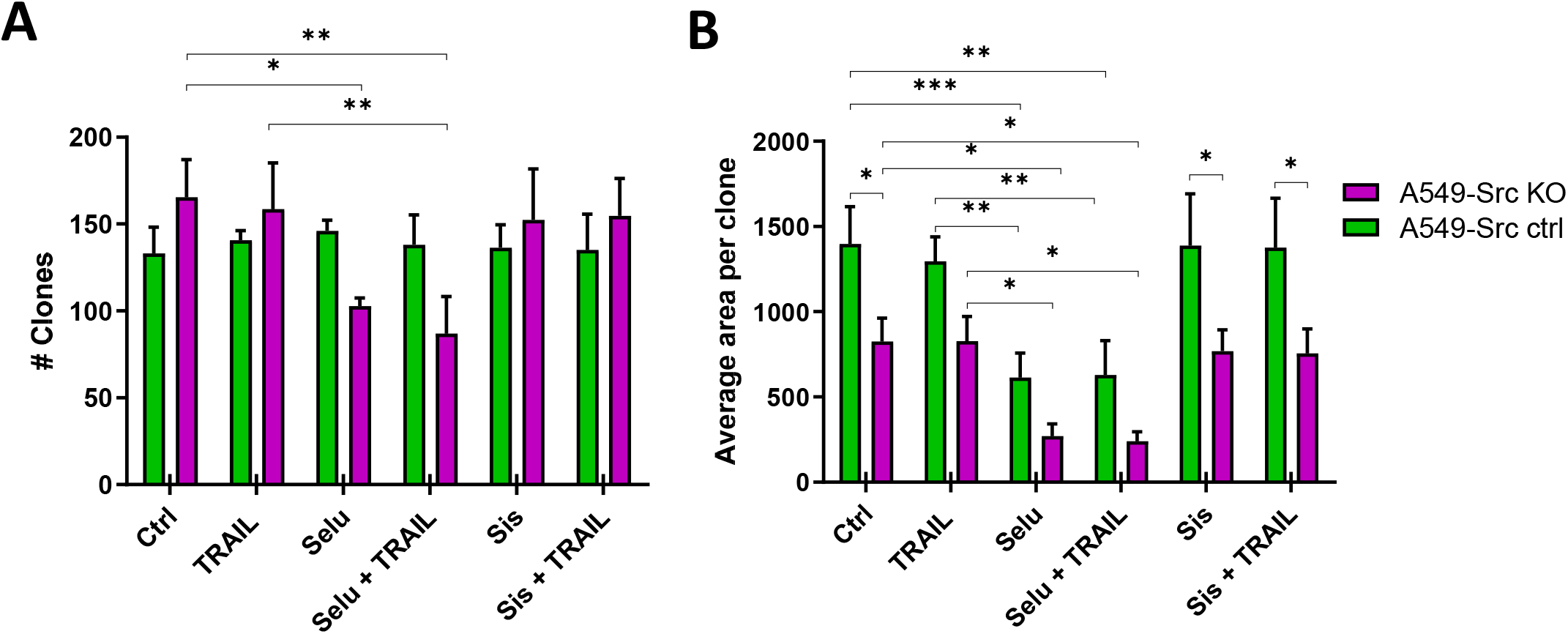
Effects of TRAIL, Src, MEK1/2 and SMAD3 on A549 in clonogenic assays. Clonogenic assays of A549-Src ctrl and -Src-KO cells with(out) rhTRAIL (50ng/ml), selumetinib (Selu, MEK1/2 inhibitor) and specific inhibitor of SMAD3 (3-sis). Colony numbers and size were determined after 10 days by imageJ software analyses. The bar graphs depict **(A)** the mean number of colonies (clonogenicity) and **(B)** the mean size of the colonies (cell growth). Independent of rhTRAIL, Src and MEK1/2 both stimulated colony growth, and MEK1/2 enhanced colony formation in Src KO cells (N=3 ± SD).

## Discussion

In this study we examined the possible function of Src as a regulator of TRAIL-induced apoptosis and mediator of pro-tumorigenic TRAIL signalling in NSCLC cells. We found different rhTRAIL-induced Src activation profiles in apoptosis sensitive H460 versus resistant A549 cells. The rapid rhTRAIL-dependent Src phosphorylation observed in

A549 cells required the presence of TRADD, a component of the secondary signalling complex. However, Src activity did not regulate sensitivity to rhTRAIL-induced apoptosis in these NSCLC cell lines. Using a Src co-IP/mass-spectrometry approach to find downstream effectors, we identified proteins that bind to Src upon rhTRAIL treatment of A549 cells. MEK1, MEK2, SMAD3 and β-catenin were selected for further analysis. Src activation was found to mediate MEK1/2 and subsequent ERK phosphorylation and SMAD3 ser423/425 phosphorylation, and to regulate β-catenin expression. Activation of these signalling pathways by Src was accompanied by pERK and β-catenin (peri)nuclear translocation. Finally, neither Src expression, nor rhTRAIL exposure could significantly affect colony formation of A549 cells, although the size of the colonies was reduced in Src deficient A549 cells, independent of rhTRAIL treatment. Inhibition of MEK1/2 reduced both the number and size of the colonies, also independently of rhTRAIL. Together these results imply that the identified Src downstream effectors regulate proliferative properties of A549 cells, however, these proteins could not be directly linked with rhTRAIL-induced Src-dependent pro-tumorigenic activity.

Our finding that Src activity does not regulate apoptosis sensitivity in the NSCLC cells is in contrast with several previous reports. For example, Src has been found to inhibit TRAIL induced apoptosis by phosphorylating procaspase 8 at Tyr380 that prevents its activation (24,26). In addition, Src inhibition in breast cancer, melanoma and hepatocellular carcinoma cells restored or increased sensitivity towards TRAIL induced apoptosis that was associated with increased caspase 8 and caspase 3 activity (26,33,43). Although, we have to extend our studies to a larger panel of NSCLC cells, it could be that the apoptosis modulatory function of Src is tumour type dependent. The underlying molecular mechanisms that cause the different functioning of Src in regulating TRAIL-induced apoptosis in different tumour types remain to be elucidated.

Previously we found that rhTRAIL via RIPK1, a kinase and component of the secondary complex, is a mediator for TRAIL-induced Src activation (44). Here we found that TRADD, another component of the secondary complex, is also required for rhTRAIL-dependent Src activation in A549 cells. We did not find involvement of FADD, caspase 8 or TRAF2 in Src activation. Our findings are consistent with the known function of TRADD, which is to recruit RIPK1 to the TRAIL receptors leading to RIPK1 activation and subsequent activation of non-apoptotic signalling cascades by preventing DISC formation and FADD-caspase 8 driven cell death (45).

Src is known to interact with various proteins often resulting in conformational changes and subsequent activation or inactivation of Src, thereby also affecting the ability of Src to interact with other proteins (46–48). Here, we examined the interactome of Src in A549 and H460 cells to identify possible mediators of pro-tumorigenic rhTRAIL signalling. Interestingly, the number of identified Src-binding proteins in untreated and rhTRAIL treated H460 cells was inverse to the numbers found in A549 and reflects the differences observed in rhTRAIL-induced Src activation in both NSCLC cell lines. In A549 cells the number of proteins in complex with Src increased after rhTRAIL exposure, likely contributing to pro-tumorigenic signalling, whereas the number of proteins decreased in H460 cells that may reflect lower protein expression during the apoptotic process at least in part due to caspase-dependent protein degradation. Surprisingly, in the Src interactome we found decreased binding of the Src inhibitor PP2A after rhTRAIL treatment in both sensitive and resistant NSCLC cells. PP2A has been reported to inhibit Src activation by dephosphorylating Src-Tyr416 and, as a consequence, the activation of apoptosis by reducing inhibitory caspase 8-Tyr380 phosphorylation. Furthermore, in resistant cells, TRAIL treatment has been found to induce ubiquitination of PP2A causing its degradation leading to Src-dependent caspase 8 inactivation (33). PP2A degradation could explain decreased binding of Src and PP2A in rhTRAIL treated A549 cells. However, such a mechanism cannot explain the decrease in Src-PP2A interactions in apoptosis sensitive H460 cells. The Src-PP2A interactions were reduced after 1 hr rhTRAIL treatment, a time point at which irreversible apoptosis is induced in H460 cells. Effector caspase-3 cleaves PP2A at the regulatory A subunit, through which its activity is increased and the apoptotic commitment of the cell is enhanced (49,50). Cleavage of PP2A might result in decreased interactions with Src and could explain why we found a sudden increase in Src phosphorylation upon 2 hrs rhTRAIL treatment. Yet, the exact mechanisms and the role of Src, caspase 8 (phosphorylation) and caspase 3 herein need to be further examined.

Differential analyses of the Src interactomes in H460 and A549 in the absence or presence of rhTRAIL allowed us to identify possible mediators of non-apoptotic TRAIL signalling. SMAD3, β-catenin, MEK1 and MEK2 were selected and their interactions with Src were confirmed by co-IPs, although this does not necessarily implies direct protein-protein interactions. Expression analyses provided evidence of rhTRAIL-dependent Src-MEK1/2-ERK, Src-β-catenin and Src-SMAD3 signalling (see also summarizing Fig. 8).

**Figure 8.**
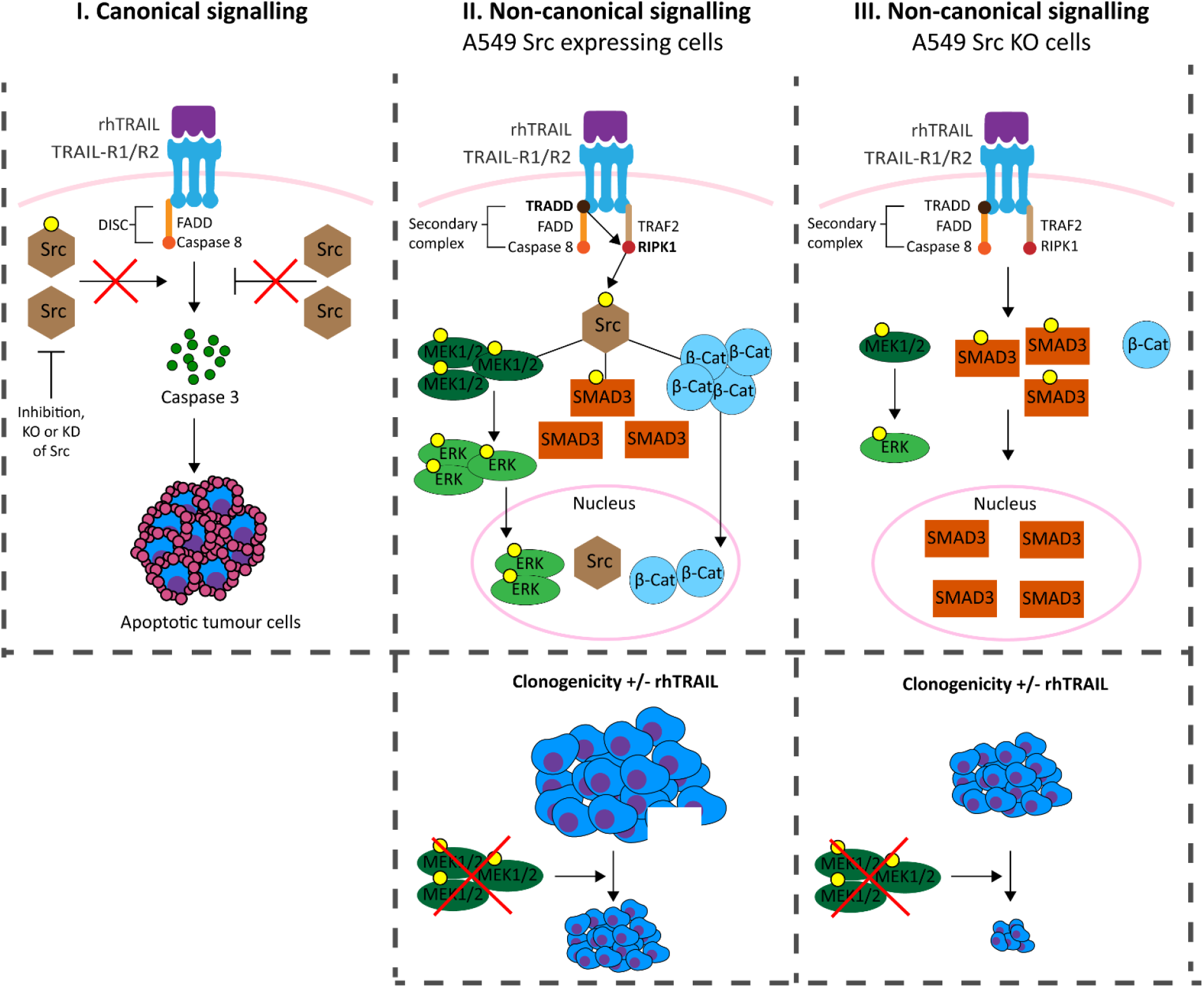
Schematic representations of the identified mechanisms by which Src can mediate TRAIL-dependent signalling involving MEK1/2-ERK, SMAD3 and β-catenin. In TRAIL resistant A549 and sensitive H460 cells **I**. Canonical signalling was not modulated by Src, as neither inhibition, knockdown nor knockout enhanced TRAIL driven cell death. In **II**. Non-canonical signalling in A549 Src expressing cells rhTRAIL activated Src by phosphorylation via TRADD-RIPK1. Src activation was persuaded by interactions with MEK1/2, SMAD3 and Catenin-β and subsequent phosphorylation of SMAD3 and MEK1/2. MEK1/2 activated ERK, which was translocated to the nucleus, as well as Catenin-β. Regarding clonogenicity, inhibition of MEK1/2 reduced the size of the clones, independent of rhTRAIL treatment. In **III**. Non-canonical signalling in A549 Src KO cells rhTRAIL induced phosphorylation of MEK1/2-ERK and SMAD3, yet only SMAD3 was translocated to the nucleus. The clones of Src deficient cells were significantly smaller in size and upon MEK1/2 inhibition their size was further reduced, as well as their number, independently of rhTRAIL exposure.

SMAD3, a downstream effector of TGF-β signalling, can have both tumour suppressive and oncogenic functions. Phosphorylation of SMAD3 at Ser423/425 has been associated with tumour suppressive activity (37,51). Interestingly, in A549 Src-KO cells rhTRAIL treatment increased pSMAD3-Ser423/425 and total SMAD3 levels in the nucleus of A549 Src-KO also increased, the latter possibly representing pSMAD3. However, pharmacological inhibition of pSMAD3 with SIS3, a selective SMAD3 inhibitor that prevents TGF-β induced phosphorylation, did not altered clonogenicity of both Src deficient and proficient A549 cells. SMAD3 is involved in TGF-β induced migration, although in our hands SMAD3 inhibition with SIS3 did not affected the migratory capacity of NSCLC (unpublished data; (52)). TRAIL can induce epithelial-mesenchymal transition (EMT) in various cancer cells, including lung tumour cells, via amongst others TGF-β/SMAD signalling pathway (53). Possibly, SMAD3 inhibition in cells undergoing EMT might demonstrate a greater impact on TRAIL non-canonical signalling, yet needs further investigation. Hence, we did not demonstrated a pro-tumorigenic function of SMAD3 in rhTRAIL-Src non-canonical signalling. β-catenin, a downstream effector of Wnt signalling, is known to promote stemness, tumorigenesis and cancer cell proliferation (38,39). Dephosphorylation of β-Catenin and subsequent protein accumulation is followed by nuclear translocation and transcriptional regulation of target genes. On the other hand, N-terminal phosphorylation of β-catenin results in ubiquitination and degradation of β-catenin. Src via activation of the focal adhesion kinase (FAK) has a pivotal role in the nuclear trans-localisation and activity of β-catenin by lowering β-catenin affinity for membrane localised E-Cadherin (54–58). rhTRAIL treatment of Src proficient cells resulted in higher basal levels of β-catenin, increased levels in the nucleus as well as at the cell membrane and peri-nuclear area. Although, we could not confirm β-catenin nuclear translocation by immunofluorescence microscopy, we found distinct spots where Src and β-catenin co-localised at the periphery of the nucleus. Src translocation to the nucleus has been previously associated with both increased and decreased tumorigenic effects in various tumour types, among which osteosarcoma, breast cancer pancreatic and acute myeloid leukaemia (59). Whether perinuclear staining implies ER localization is unclear to us, as well as the possible functional consequences of perinuclear co-localisation. Overall, the role of Src in altering the subcellular localisation of β-catenin remains to be further elucidated.

MEK1 and MEK2 are part of the RAS mediated RAF-MEK1/2-ERK proliferation and survival pathway that is often hyperactivated in lung cancers (35,36). We found that depletion of Src and simultaneous inhibition of MEK1/2 reduced the clonogenicity of rhTRAIL resistant A549 cells, which was stronger than after MEK1/2 inhibition or Src ablation alone. Simultaneous treatment with pharmacological inhibitors of Src and MEK in NSCLC and ovarian cancer *in vitro* and *in vivo* has shown synergistic anti-tumour effects (60–62). Further studies are needed to confirm and examine whether and how these proteins are involved in TRAIL-Src non-canonical signalling in resistant NSCLC cells.

Taken together, our results provide deeper insight in the possible mechanisms underlying TRAIL-RIPK1-Src mediated pro-tumorigenic signalling. Src was not instrumental for causing apoptosis resistance in A549 cells, but could modulate MEK1/2, SMAD3 and β-catenin signalling, although their involvement in pro-tumorigenic signalling should be further corroborated. Whether these possible pro-tumorigenic pathways provide therapeutic targets that would increase the efficacy of TRAIL in resistant NSCLC cells remain to be investigated.

## Acknowledgements

CRIPSR/Cas knock outs were generated with help from the iPSC/CRISPR Centre, ERIBA, UMCG, University of Groningen. Mass spectrometry was performed with the help of M.P. de Vries of the Interfaculty Mass Spectrometry Center Groningen, Laboratory of Paediatrics, UMCG, University of Groningen. We also thank Vincent Leeuwenburgh for his help with graphical displays of the mass-spec data.

