## Supplemental Figures for "TRAIL-induces Src mediated MEK/ERK, SMAD3 and β-catenin signalling in apoptosis resistant NSCLC cells"

### Supplementary Figures

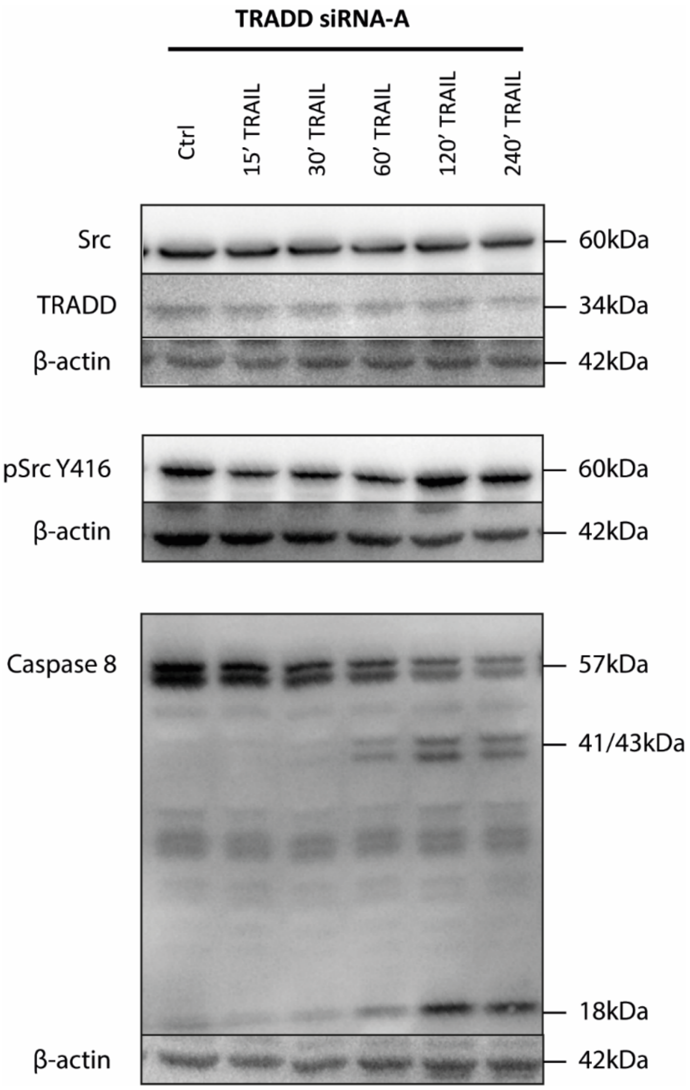


**Supplementary Figure 1. TRADD2 is required for early Src phosphorylation after rhTRAIL exposure**

Western blot analysis depicting Src, TRADD, Src-Y418p and caspase 8 protein levels in A549 cells in absence or presence of rhTRAIL (50ng/ml) after shRNA-mediated silencing of TRADD, using a different siRNA than used in Fig. 2.

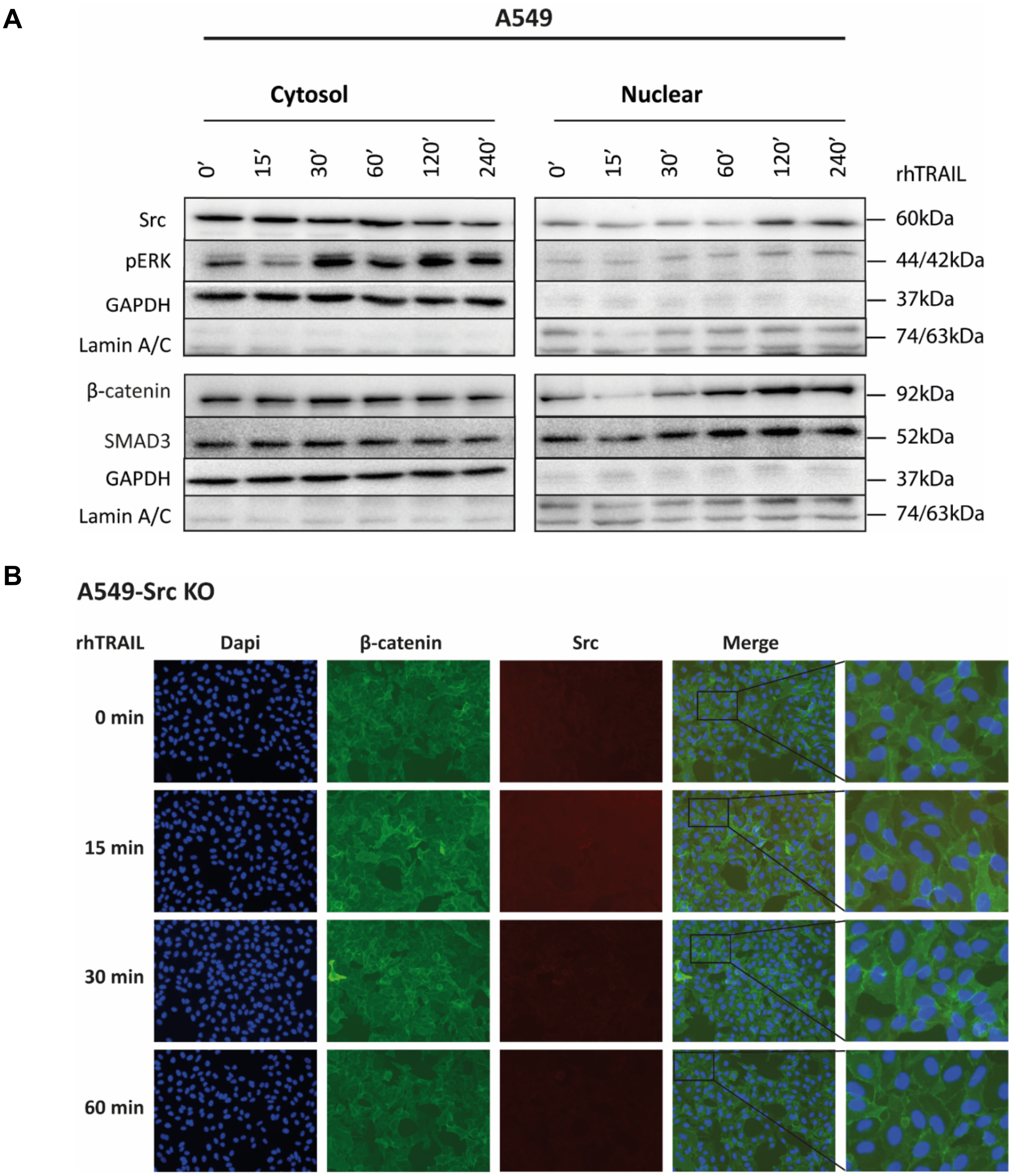


**Supplementary Figure 2. Subcellular localisation of Src, MEK1, MEK2, SMAD3 and β-catenin after rhTRAIL treatment**

**(A)** Western blot analysis showing Src, MEK1, MEK2, SMAD3 and β-catenin protein levels in the cytosol and nucleus from A549 cells after treatment with rhTRAIL (50 ng/ml) for the indicated time periods. Expression patterns were similar as seen in A549-Src ctrl cells (Fig 6B). **(B)** Immunofluorescent microscopy of A549 Src-KO cells stained with DAPI (blue), β-catenin (green) and Src (red) and merged pictures with enlargements. Staining of Src and β-catenin were determined after 15, 30 and 60 minutes rhTRAIL treatment. There was no increased β-catenin localisation at the plasma membrane and peri-nuclear area as we found for A549-Src control cells (Fig 6C).
